# Reduced Proactive and Reactive Cognitive Flexibility in Older Adults Underlies Performance Costs During Dual-Task Walking: A Mobile Brain/Body Imaging (MoBI) Study

**DOI:** 10.1101/2024.01.27.577090

**Authors:** David P. Richardson, John J. Foxe, Edward G. Freedman

## Abstract

Age-related reductions in cognitive flexibility may limit modulation of control processes during systematic increases to cognitive-motor demands, exacerbating dual-task costs. In this study, behavioral and neurophysiologic changes to proactive and reactive control during progressive cognitive-motor demands were compared across older and younger adults to explore the basis for age-differences in cognitive-motor interference (CMI). 19 younger (19 - 29 years old, mean age = 22.84 +/- 2.75 years, 6 male, 13 female) and 18 older (60 - 77 years old, mean age = 67.89 +/- 4.60 years, 9 male, 9 female) healthy adults completed cued task-switching while alternating between sitting and walking on a treadmill. Gait kinematics, task performance measures, and brain activity were recorded using electroencephalography (EEG) based Mobile Brain/Body Imaging (MoBI). Response accuracy on easier trial types improved in younger, but not older adults when they walked while performing the cognitive task. As difficulty increased, walking provoked accuracy costs in older, but not younger adults. Both groups registered faster responses and reduced gait variability during dual-task walking. Older adults exhibited lower amplitude modulations of proactive and reactive neural activity as cognitive-motor demands systematically increased, which may reflect reduced flexibility for progressive preparatory and reactive adjustments over behavioral control.

## INTRODUCTION

Maintaining gait stability and cognitive task performance during dual-task walking requires adjustments to cognitive control (Hausdorff, Schweiger, Herman, Yogev-Seligmann, & Giladi, 2008; Herman, Mirelman, Giladi, Schweiger, & Hausdorff, 2010). Gait consistency may improve when the simultaneous cognitive demands are mild (Beilock, Carr, MacMahon, & Starkes, 2002; Beilock & Gray, 2012; Hamacher, Koch, Löwe, & Zech, 2019; Huxhold, Li, Schmiedek, & Lindenberger, 2006; Lövdén, Schaefer, Pohlmeyer, & Lindenberger, 2008; Patelaki, Foxe, Mazurek, & Freedman, 2022; Richardson, Foxe, Mazurek, Abraham, & Freedman, 2022), but dual-task costs to gait integrity and cognitive performance often arise when sensorimotor and cognitive demands coincide (Al-Yahya et al., 2011; Hausdorff et al., 2008; Huxhold et al., 2006; Plummer et al., 2013; Sheridan, Solomont, Kowall, & Hausdorff, 2003; Springer et al., 2006; Woollacott & Shumway-Cook, 2002). Reproducible associations between dual-task costs, advancing age, and pathology have motivated the development of dual-task evaluations of fall risk and cognitive health (Hausdorff et al., 2008; Herman et al., 2010; Li, Bherer, Mirelman, Maidan, & Hausdorff, 2018; M. Montero-Odasso, Muir, & Speechley, 2012; M. Montero-Odasso, Verghese, Beauchet, & Hausdorff, 2012; M. M. Montero-Odasso et al., 2017; Muir, Gopaul, & Montero Odasso, 2012; Sheridan & Hausdorff, 2007; Springer et al., 2006). The neurophysiology underlying dual-task cost progression with aging has not been thoroughly characterized. Establishing neurophysiological markers of age- and pathology- related changes in dual-task performance can enrich our understanding of degenerative processes, increase capacity for early detection of affected or at-risk individuals, and provide therapeutic benchmarks for novel cognitive-motor training regimens (e.g., (Lizarraga et al., 2023)).

Cognitive flexibility underlies adaptation to overt shifts in demands and priorities (Diamond, 2013; Miyake et al., 2000). Prefrontal cortex (PFC) is recruited when automatic processes within cognitive, sensory and motor domains are insufficient or inappropriate, and a shift to active control is required (Braver, 2012; Braver, Paxton, Locke, & Barch, 2009; Cole, Bassett, Power, Braver, & Petersen, 2014; De Sanctis, Gomez-Ramirez, Sehatpour, Wylie, & Foxe, 2009; Menon & D’Esposito, 2022; Paxton, Barch, Racine, & Braver, 2008). These processes can be deployed proactively, in anticipation of impending demands, or reactively, on an ‘as needed’ basis during the course of a demanding event (Braver et al., 2009; Foxe & Simpson, 2005; Foxe, Simpson, Ahlfors, & Saron, 2005; Wylie, Javitt, & Foxe, 2003a, 2006). As the brain ages, reliance upon frontal cortical structures increases as sensory, cognitive, and motor domains are degraded by age-related cortical atrophy (Fettrow et al., 2021; Holtzer, Epstein, Mahoney, Izzetoglu, & Blumen, 2014; Mirelman et al., 2017; Park & Reuter-Lorenz, 2009).

Capacity to compensate may be limited by the anterior to posterior sequence of cortical atrophy, during which PFC grey matter volume reduction and frontal white matter atrophy emerge as early structural signs of neurocognitive aging (Lemaitre et al., 2012). Functional consequences of neurocognitive aging on control networks include reduced proactive control, and a resultant shift towards reactive control modes (Braver, 2012; Gajewski, Ferdinand, Kray, & Falkenstein, 2018; Jimura & Braver, 2009; Kopp, Lange, Howe, & Wessel, 2014; Paxton et al., 2008). Together, these changes suggest aging brains increasingly require effortful cognitive control over behaviors, but with diminished capacity to provide that control. These age-related changes are often sufficient to degrade performance when only a single task is performed. Overlap in demand for cognitive control during dual-task walking may further stress network vulnerabilities beyond the limits for adaptation.

The underlying theme of insufficient supply for demands which are too great or urgent is often addressed with brain structure models of dual-task costs. The central capacity limitation model conceives dual-task costs as outcomes of processing shortfalls when demands exceed structural limits on cognitive control (Friedman, Polson, Dafoe, & Gaskill, 1982; Kahneman, 1973; Koch, Poljac, Muller, & Kiesel, 2018; Tombu & Jolicœur, 2003). Bottleneck models interpret dual-task costs as the outcome of a “first-come-first-served” competition over control network elements which are structurally unable to sustain task processes in parallel (Koch et al., 2018; Pashler, 1994; Ruthruff, Pashler, & Klaassen, 2001; Wylie, Javitt, & Foxe, 2004a). These models are not mutually exclusive, and suggest that higher dual-task costs in older adults stem from structural changes and overall brain mass reduction which diminish capacity limits or narrow bottleneck points in control networks. This aligns with reported correlations between dual-task costs, and structural preservation of PFC, medial frontal gyrus, and cingulate cortex during aging (Allali et al., 2019; Hupfeld et al., 2022; Tripathi, Verghese, & Blumen, 2019; Wagshul, Lucas, Ye, Izzetoglu, & Holtzer, 2019). Older and younger adults do not appear to share identical links between brain structure and dual-task walking performance (Hupfeld et al., 2022). Age-specific structural-functional relationships could reflect the changing demand for cortical recruitment during cognitive-motor tasks across the lifespan, but functional neurophysiological measures during active motion are required to elucidate these differences.

Cued task-switching is a prototypical paradigm for testing cognitive flexibility through comparison of proactive and reactive control processes across trials, both with and without attention switching requirements. Three types of trials comprise the typical task-switching paradigm: pure trials (a single task is repeated in a block *without* switching requirements), mixed repeat trials (a task is immediately repeated in a sequence of trials during blocks of stimuli *with* switching requirements), and switch trials (the task of the preceding trial ceases, and a different task is performed) (Allport, Styles, & Hsieh, 1994; Foxe, Murphy, & De Sanctis, 2014; Monsell, 2003; Weaver, Foxe, Shpaner, & Wylie, 2014; Wylie, Javitt, & Foxe, 2003b; Wylie, Murray, Javitt, & Foxe, 2009). Proactive and reactive neural activity of younger adults demonstrates larger amplitude modulations across attention switching requirements than that of older adults. The cue-evoked centroparietal positivity and frontal negativity are two preparatory signals associated with the posterior parietal cortex (PPC) and dorsolateral prefrontal cortex (DLPFC) respectively (S. Jamadar, Hughes, Fulham, Michie, & Karayanidis, 2010; Frini Karayanidis & Jamadar, 2014; F. Karayanidis et al., 2009). Both components are amplified in anticipation of switch trials in younger adults, but not older adults (Gajewski & Falkenstein, 2011; Gajewski et al., 2018; Gajewski et al., 2010; F. Karayanidis, Whitson, Heathcote, & Michie, 2011). Target-evoked P2, N2, and P3 are three reactive signals associated with activity in the anterior cingulate cortex (ACC), supplementary motor area (SMA), and PPC (Iannaccone et al., 2015; Irlbacher, Kraft, Kehrer, & Brandt, 2014; S. D. Jamadar, Thienel, & Karayanidis, 2015). In younger adults, both the P2 and target-P3 are attenuated on switch trials, but the N2 is amplified. In older adults, these switch-specific changes to reactive control processes are again reduced (S. D. Jamadar et al., 2015; Frini Karayanidis & Jamadar, 2014; F. Karayanidis, Whitson, et al., 2011; Kieffaber & Hetrick, 2005; Mansfield, Karayanidis, & Cohen, 2012; Whitson et al., 2014). Dual-task walking further accentuates these proactive and reactive processes in younger adults (Richardson et al., 2022). Older adults, on the other hand, generally show lower amplitude modulations of anticipatory and reactive neural signals across pure, mixed repeat, and switch trials (Gajewski et al., 2018).

Functional models for dual-task walking costs focus on the capacity to flexibly adapt control processes to shifting cognitive and motor demands. EEG-based Mobile Brain/Body Imaging (MoBI) techniques permit comparison of functional neurophysiological activity between single and dual-task walking conditions with excellent temporal resolution and an established record of long-term test-retest reliability (De Sanctis, Butler, Green, Snyder, & Foxe, 2012; De Sanctis, Butler, Malcolm, & Foxe, 2014; De Sanctis et al., 2020; De Sanctis et al., 2023; Gramann, Ferris, Gwin, & Makeig, 2014; Gramann, Gwin, Bigdely-Shamlo, Ferris, & Makeig, 2010; Gramann et al., 2011; Gramann, Jung, Ferris, Lin, & Makeig, 2014; Makeig, Gramann, Jung, Sejnowski, & Poizner, 2009; Malcolm, Foxe, Butler, & De Sanctis, 2015; Malcolm, Foxe, Butler, Molholm, & De Sanctis, 2018; Malcolm et al., 2019; Patelaki et al., 2022; Richardson et al., 2022). When performing a single task, proactive control processes bring individuals to a state of readiness in anticipation of demanding events (Dale, Simpson, Foxe, Luks, & Worden, 2008; Kelly, Gomez-Ramirez, & Foxe, 2009; Posner & Petersen, 1990). Reactive processes subsequently coordinate execution of the task and corrective responses to unexpected interference (Braver, 2012; Braver et al., 2009; Jimura & Braver, 2009; Morie, De Sanctis, Garavan, & Foxe, 2016). Age-related, environmental, and experimental disruptions to the integrity of either proactive or reactive cognitive control results in performance decline (P. S. Cooper, Garrett, Rennie, & Karayanidis, 2015; Patrick S. Cooper et al., 2019; P. S. Cooper, Wong, et al., 2015; S. D. Jamadar et al., 2015; Frini Karayanidis & Jamadar, 2014; F. Karayanidis, Whitson, et al., 2011; Kopp et al., 2014; Mansfield et al., 2012; Whitson et al., 2014). Controlling walking-related disturbances to proactive and reactive control processes is crucial to maintaining cognitive performance during dual-task walking.

Event-related potential (ERP) indices of both proactive and reactive cognitive control are progressively accentuated by walking in healthy younger adults (Richardson et al., 2022). In a previous MoBI study comparing younger and older adult neural responses during dual-task walking, accentuation of inhibitory control processes was attenuated and occurred at later processing stages in older adults (Malcolm et al., 2015). The older adults also suffered dual-task costs to performance, whereas younger adults did not. Delayed and diminished accentuation of control processes in the context of larger dual-task performance costs were interpreted as neurophysiological indicators of reduced flexibility in the control processes of older adults. Lower amplitude accentuation during dual-task conditions may be a feature of reduced adaptation to cognitive-motor demands irrespective of age (Patelaki, Foxe, Mantel, Kassis, & Freedman, 2023; Patelaki et al., 2022). Among the likely many factors which contribute to successful cognitive-motor adaptation, preparation in advance of demanding cognitive events is characteristically reduced within the reactive modes of control which predominate in older adults (Braver, 2012; Gajewski et al., 2018; Kopp et al., 2014). Reactive modes of control may neglect opportunities for proactive cognitive-motor adaptation, and subsequently shift greater cognitive-motor loads to clash with critical reactive processes. We hypothesize age-related reductions to flexibility of proactive and reactive control processes limit adaptation to cognitive-motor demands, worsening dual-task costs. To test this hypothesis, we use a cued task-switching paradigm to compare neural activity evoked by trials with and without attention switching requirements. We predicted that dual-task walking would not accentuate proactive and reactive processes in older adults to the degree observed in younger adults, and we expected dual-task costs to gait and cognitive performance to be higher. Larger amplitude proactive and reactive neural activity across cognitive-motor demands may reflect control networks capable of greater dynamic range, and be a marker of superior capacity for adapting one’s behavior to dual-task walking without performance degradation.

## MATERIALS AND METHODS

### Participants

19 young adults (19 - 29 years old, mean age = 22.84 +/- 2.75 years, 6 male, 13 female) and 18 older adults (60 - 77 years old, mean age = 67.89 +/- 4.60 years, 9 male, 9 female) completed the cued task-switching experiment, providing the data analyzed in this report. All participants included are original to the present dataset, and have not been included in any previously published datasets. All participants provided written informed consent, reported no diagnosed neurological conditions, no recent head injuries, normal or corrected-to-normal vision, and that they were not suffering post-concussive syndrome. All procedures were approved by the University of Rochester Institutional Review Board (STUDY00001952), and complied with the ethical principles of the Declaration of Helsinki for the responsible conduct of research. Participants were paid $18/hour for time spent in the lab.

### Stimuli and Task

We employed a so-called “cued task-switching” paradigm, a schematic of which is depicted in **Figure 1** and which has been described in detail in our prior work (Richardson et al., 2022). Participants were presented with a central cue stimulus lasting 100ms that instructed them which of two tasks they would be required to perform. At an interval of 650ms following the cue offset, a second 100ms duration stimulus set appeared, which consisted of bilateral gabor patch images. Participants used handheld devices to select the gabor patch with either higher spatial frequency, or further clockwise orientation, depending on which task they were performing (Fig. 1A). That is, the same gabor patch stimuli afforded two different tasks – one a judgement based on spatial frequency, and the other a judgment based on the orientation of the gratings within the patch. In “pure blocks” participants repeatedly performed the same task on every trial (Fig. 1B). In “mixed blocks”, participants would switch between the tasks when the cue changed (Fig. 1C). A central-fixation cross was held on-screen continuously throughout experimental blocks. Participants performed the task while sitting, and at other times while walking on a treadmill (Tuff Tread, Conroe, TX, USA). The specific features of the cue and task stimuli are described in detail in what follows.

**Figure 1:**
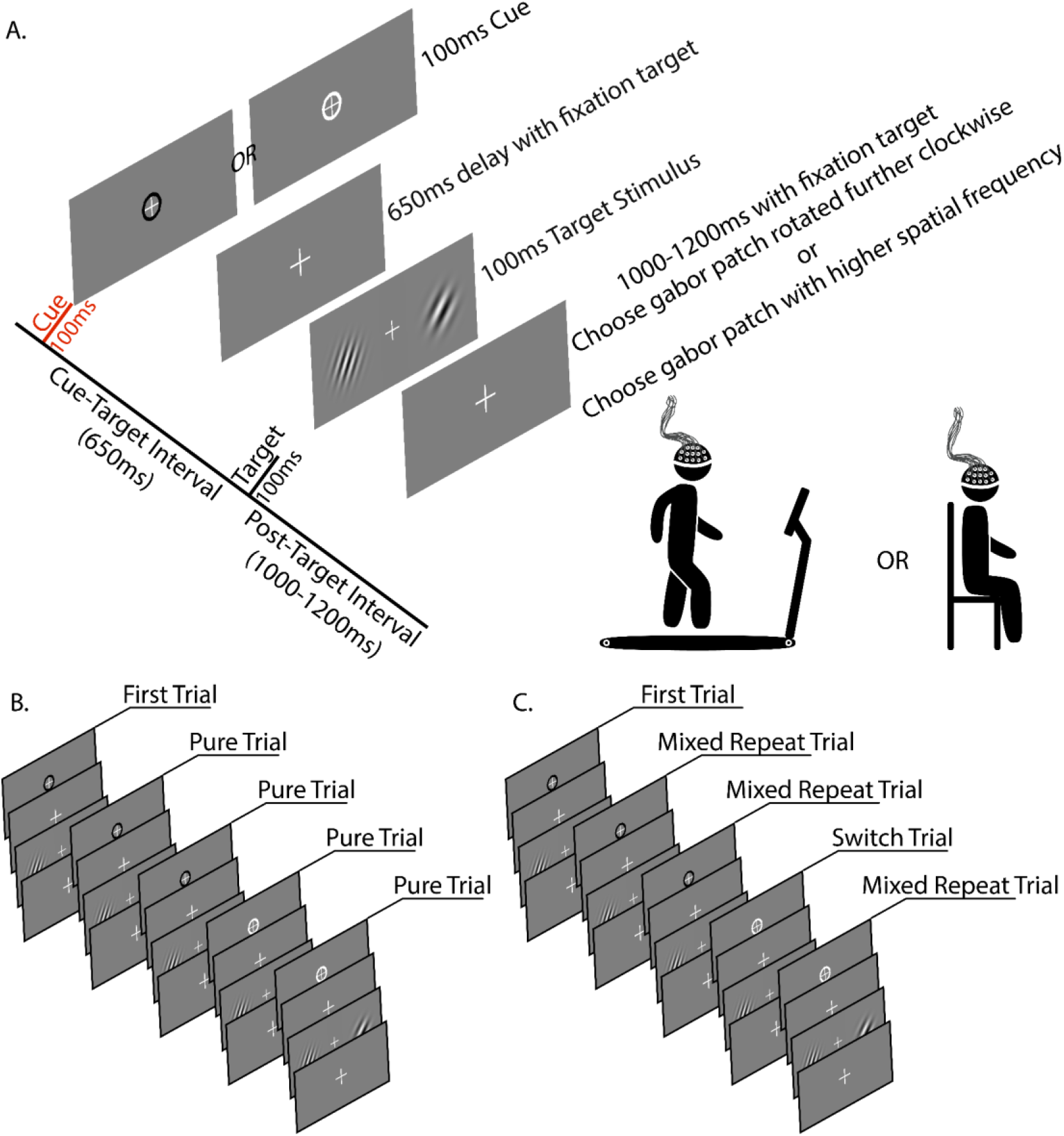
Experimental procedure overview. **(A)** Description of trial elements, timing, and duration. Participants walked on a treadmill or sat in a pseudorandom order**. (B)** Example trial sequence of a Pure Block. **(C)** Example trial sequence of a Mixed Block. Switch trials are signaled by a change in cue color. Note the first trial in each block is excluded from analyses as it is neither a Repeat, nor a Switch trial.

All visual task elements were projected on a screen 2.2m (during seated blocks) in front of the participant. A black or white annulus encircling the central-fixation cross served as a cue on each trial, instructing participants to perform either the spatial frequency discrimination or the orientation discrimination task. Target stimulus sets comprised two bivalent gabor patches (differing in both orientation, and spatial frequency) presented to the left and right of the central fixation cross. Stimulus sets were selected from 200 possible pairings to ensure incongruent task response mappings to the left- and right-hand response devices (Nintendo Switch Joy-Con^™^ controllers). “Replace trials” consisting of identical stimuli (i.e., the right and left patches were the same) were presented intermittently throughout the experiment (4% of trials). Based on the location of the treadmill, eccentricity to the near edge, center, and far edge of each gabor patch was 4.27°, 6.40°, and 8.50° respectively left and right of the central fixation during sitting blocks.

Experimental blocks consisted of 150 trials. Pure blocks required participants to repeatedly perform either the orientation task or the spatial frequency task, but never both in the same block (i.e., no task switching). In mixed blocks, participants switched between tasks when instructed by a change in cue color. Each trial consisted of a 100ms cue, a 650ms cue-target interval (CTI), a 100ms target stimulus set, and a 1000-1200ms post target interval (PTI), after which the next trial commenced immediately. Cues were delivered during both pure blocks (non-informative) and mixed blocks (informative). Cues would repeat a minimum of 4 times, after which the probability of a cue switch would increase on each subsequent trial. A maximum of 6 cue repetitions (1 switch trial + 5 repeats) were permitted in mixed blocks. A minimum of 4 pure blocks were administered, 2 seated, 2 walking. A minimum of 10 mixed blocks were administered (5 seated, 5 walking). Walking only blocks (walking without performing the cognitive task) were also administered at pseudorandom intervals. Participants were instructed to continue fixating on the central fixation cross during walking only blocks, but no stimuli were presented, and subjects did not perform a cognitive task. Walking and sitting block order was pseudorandom. No more than 3 consecutive sitting or walking blocks could occur.

### Procedure

Participants and research personnel collaboratively determined each participants natural walking pace at the beginning of the experiment (younger adult mean speed = 1.17 m/s +/- 0.1185; older adult mean speed = 0.70 m/s +/- 0.2127). The same walking speed was used throughout the experiment. Experimental blocks lasted 4-6 minutes. Short breaks occurred between every block (1-3 minutes) during which snacks and water were made available to the participant. Longer breaks could be taken at the participant’s request. Each experiment was conducted in a single session lasting fewer than 5 hours.

Participants were guided through a training block with target duration increased to 8 seconds to permit verbal feedback from research coordinators on selected responses. Response feedback was not provided during the recorded experimental blocks. The instructions provided for the orientation discrimination task were to select the gabor patch that was rotated further clockwise using the response devices in the left or right hand to indicate the left or right side target respectively. Research personnel informed participants that gabor patches with vertical lines were not rotated at all, horizontal lines represented 90 degrees of rotation, and target stimuli would only be rotated between 0 and 90 degrees. In the spatial frequency discrimination task, participants were instructed to select the gabor patch with higher spatial frequency (i.e., more/denser lines). During pure blocks, participants were instructed that the cue color did not provide performance feedback, or any prospective information beyond indicating a target stimulus was imminent. Cue color-task mappings were revealed prior to the first mixed block, and were counterbalanced across participants in both age groups. Research personnel actively confirmed participant comprehension of cue-task mappings during breaks throughout the remainder of the experiment.

### Cued Task-Switching Performance Analysis

The sensitivity index (d’) and response times for trials performed in pure and mixed blocks were compared to quantify costs associated with resolving interference from multiple tasks during mixed blocks (mixing costs) (Green & Swets, 1966; Monsell, 2003; Rogers & Monsell, 1995; Shaw et al., 2020; Wylie et al., 2009). The costs related to actually switching between tasks within mixed blocks (switch costs) were assessed by comparing mixed repeat and switch trials (Wylie, Javitt, & Foxe, 2004b). Responses were considered valid if they occurred in the window 200-1000ms following target stimulus onset. Trials in which no response was provided were excluded from analyses. D-prime (d’), a signal detection measure, was used to assess response accuracy and calculated according to the following equation:

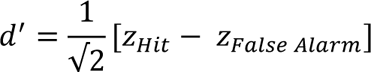

In two-alternative forced choice experiments, the classification of “hits” and “false alarms” is arbitrary (Macmillan & Creelman, 2004). Here, “hits” were defined as trials in which the “correct” target stimulus image fell on the left side and the left response device was selected. “False alarms” were defined as trials in which the “correct” target stimulus image fell on the right side, but the left response device was selected. Altering these classifications does not change resultant d’ values. Mixing costs and switching costs were assessed using two separate 2×2×2 repeated measures ANOVAs with factors of trial type (pure and mixed repeat, or mixed repeat and switch), physical state (sitting or walking), and age group (young or old). Participant d’ values and mean response times were used as dependent measures.

Tukey’s honest significant differences (HSD) test was used for *post hoc* analyses of switch costs. Significance threshold was set at α = 0.05. Dual-task effects on performance were categorized into “improvement”, “no change”, or “cost” according to the procedure used by Patelaki and colleagues (Patelaki, Foxe, Mantel, et al., 2023; Patelaki et al., 2022). Walking-minus-sitting d’ values and response times for each participant were defined as dual-task improvement or cost if they fell outside the 95% confidence interval of the normal distribution centered around a mean of zero and with a standard deviation equal to the variance of the entire cohort of younger and older adults.

### Gait Recording and Analysis

Kinematic data were recorded at 360 Hz using an optical motion capture system (Optitrack Prime 41 cameras, Motive v2.1 software, Natural Point Corvallis, OR, USA). 41 reflective markers were placed over anatomic landmarks on motion capture suits worn by participants. 4 markers on each foot (posterior of the calcaneus, tip of the second distal phalanx, distal end of the first metatarsal, and distal end of the fifth metatarsal) were used to measure stride length, stride length variability, stride time, and stride time variability.

Processing and analysis of kinematic data were performed using MATLAB 2019b (Mathworks, Natick, MA, USA). Motion capture data from each walking block were lowpass filtered at 6 Hz with a 2nd order, zero-phase Butterworth filter. Intervals of marker occlusion were interpolated. Strides were excluded from analysis if a heel marker was occluded for longer than 1 gait cycle. The filtered signal was copied, and filtered again using a 25-point moving average to smooth remaining artifact. Automatic peak detection identified extreme horizontal values coincident with vertical movement onset and offset in the antero- posterior plane. Resultant points were classified as heel strikes and lifts in the 6 Hz lowpass filtered data. Review of motion capture confirmed timing of heel strikes and lifts.

One younger adult was excluded from gait analysis due to equipment failure. Walking data from the remaining 18 younger and 18 older participants were classified as walking only, walking during a pure block, or walking during a mixed block. A minimum of 200 consecutive strides was required for a block to be included in analysis. Stride length was extracted by summing the horizontal distance between opposite heel markers at heel lifts and subsequent heel strikes. Stride time was measured as time elapsed between two consecutive heel strikes by the same foot. Coefficient of variation (CV% = (SD/mean) × 100) was used for measures of stride variability (Hausdroff, 2005). Normalized kinematic values were calculated for strides in pure and mixed blocks relative to the baseline gait during walking only blocks when no cognitive task was being performed according to the following equation:

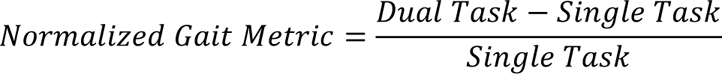

Negative normalized values indicated a reduction in stride length, time, or variability during dual-task walking. 3×2 repeated measures ANOVAs with factors of block type (walking only, pure, mixed) and age group (younger or older adults) compared mean stride length, stride time, stride length variability, and stride time variability. Results are presented in Table 2. When appropriate, Greenhouse-Geisser correction was used to account for violation of the assumption of sphericity. Tukey’s HSD was used for *post-hoc* analysis and effect sizes were calculated using Cohen’s d.

### EEG Recording and Pre-Processing

Eelectroencephalographic (EEG) data were recorded from 128 scalp channels and continuously digitized at 2048 Hz throughout the duration of the experiment (BioSemi ActiveTwo, Amsterdam, The Netherlands) and referenced to the Common Mode Sense (CMS) and Driven Right Leg (DRL). Signals were streamed via LabStreamingLayer (LSL) to a computer along with task events, and motion capture data. EEG data were visually inspected, and data corresponding to block intermissions were removed. Signals were bandpass filtered 1-50 Hz, and bad channels were identified on the basis of being flat for > 5s, and/or correlation to nearest neighbors < 0.8. Bad channels were removed and interpolated. Next, data were down-sampled to 256 Hz, datasets were re-referenced to the average reference, and adaptive mixture ICA (Palmer, Makeig, Kreutz-Delgado, & Rao, 2008; Sejnowski, 1996) was performed. Resultant ICs were inspected, and classified into 7 categories (Brain, Eye, Muscle, Heart, Channel Noise, Line Noise, Other) using an automated component classifier (ICLabel v1.2.5) (Pion-Tonachini, Kreutz-Delgado, & Makeig, 2019) that computes IC class probabilities for components. ICs with a brain classification below 0.7 were excluded from analysis. Remaining ICs were visually inspected for violation of the expected 1/f power spectral density, absence of a scalp topography consistent with an intracranial generator, and high-variance noise matching the incidence of heel strikes. ICs were back-projected onto continuous data, maintaining the same band-pass filter settings (1-50 Hz). Cue-locked epochs spanning the time-window from 100ms preceding cue onset to 1200ms after target stimulus onset were extracted relative to a 100ms pre-cue baseline. Trials were rejected on the basis of +/-85µV amplitude and 3 standard deviation threshold limits. Only trials responded to correctly 200- 1000ms after target onset were included in subsequent analyses. EEG preprocessing was performed using the EEGLAB toolbox (EEGLAB v14.1.2b) (Delorme & Makeig, 2004) and custom MATLAB scripts. After preprocessing, an average of (mean per subject +/- 1 SD) 198.08 +/- 35.23 sitting pure trials, 373.94 +/- 67.98 sitting mixed repeat trials, 95.61 +/- 18.59 sitting switch trials, 190.17 +/- 40.56 walking pure trials, 357.83 +/- 66.01 walking mixed repeat trials, and 92.58 +/- 20.32 walking switch trials remained for analysis.

### ERP Data analysis

Channel clusters at four preplanned time windows extending from 500-750ms (CTI), 900-950ms (PTI), 1000-1150ms (PTI), and 1350-1550ms (PTI) were defined by characterizing spatial and temporal trends in cluster plots of statistically significant differences between the sitting and walking neural responses from a previously published dataset of young adults performing the same task (Richardson et al., 2022). After identifying the bounds of statistically significant differences, channel clusters and time intervals were further trimmed to focus on areas of maximal differences using topographical maps of walking-*minus*-sitting differences. Mean amplitude of the channel cluster grand average across the resultant time intervals was used as the dependent measure in subsequent analyses. 2×3×3×2 repeated measures ANOVAs with factors of physical state (sitting or walking), trial type (pure, mixed repeat, or switch), channel cluster, and age group (younger or older adults) were conducted at the 500-750ms, 1000-1150ms, and 1350-1550ms intervals. A 2×3×2 repeated measures ANOVA was performed at the 900-950ms interval as there was only a single channel cluster. Greenhouse-Geisser correction was used to account for violation of the assumption of sphericity when appropriate, and effect sizes were calculated using partial eta squared (η²_p_) (Table 3). Tukey’s honest significant differences (HSD) tests were conducted for post-hoc analyses, and effect sizes were calculated using Cohen’s d. Significance threshold was set at α = 0.05.

## RESULTS

To improve readability of the text, only *p*-values for main and interaction effects of the ANOVAs are presented in the following sections. Corresponding F-statistics and effect sizes (η²_p_) are organized in Tables 1 (task performance), 2 (kinematics), and 3 (ERP measures). Both *p*-values and effect sizes (Cohen’s d) are presented within the text for post-hoc comparisons.

**Table 1:** Results of repeated measures ANOVAs comparing d’ and RT to assess age and walking- related effects on mixing costs (pure vs. mixed repeat trials) and switch costs (mixed repeat vs. switch trials). Bold text indicates *p*-value < 0.05. Effect size for ANOVA is reported as partial eta squared (η²_p_).

|  | Mixing Cost |  | Switch Cost |  |
| --- | --- | --- | --- | --- |
|  | D-Prime | Response Time | D-Prime | Response Time |
| Physical State | **F(1,35)=4.345**<br>**p=0.045**<br>**$\eta^2_p=0.110$** | **F(1,35)=17.79**<br>**p=1.655e-04**<br>**$\eta^2_p=0.337$** | F(1,35)=0.260<br>p=0.613<br>$\eta^2_p=0.007$ | **F(1,35)=18.779**<br>**p=1.178e-04**<br>**$\eta^2_p=0.349$** |
| Trial | **F(1,35)=31.926**<br>**p=2.232e-06**<br>**$\eta^2_p=0.477$** | **F(1,35)=10.393**<br>**p=0.003**<br>**$\eta^2_p=0.229$** | **F(1,35)=39.22**<br>**p=3.484e-07**<br>**$\eta^2_p=0.528$** | **F(1,35)=14.077**<br>**p=6.357e-04**<br>**$\eta^2_p=0.287$** |
| Group | F(1,35)=1.272<br>p=0.267<br>$\eta^2_p=0.035$ | **F(1,35)=16.627**<br>**p=2.492e-04**<br>**$\eta^2_p=0.322$** | F(1,35)=0.399<br>p=0.532<br>$\eta^2_p=0.011$ | **F(1,35)=19.609**<br>**p=8.898e-05**<br>**$\eta^2_p=0.359$** |
| Physical State<br>* Group | **F(1,35)=4.249**<br>**p=0.047**<br>**$\eta^2_p=0.108$** | F(1,35)=0.907<br>p=0.347<br>$\eta^2_p=0.025$ | **F(1,35)=7.535**<br>**p=0.010**<br>**$\eta^2_p=0.177$** | F(1,35)=0.279<br>p=0.601<br>$\eta^2_p=0.008$ |
| Trial * Group | F(1,35)=2.52<br>p=0.121<br>$\eta^2_p=0.071$ | F(1,35)=0.415<br>p=0.524<br>$\eta^2_p=0.012$ | F(1,35)=0.106<br>p=0.746<br>$\eta^2_p=0.003$ | F(1,35)=0.885<br>p=0.353<br>$\eta^2_p=0.025$ |
| Physical State<br>* Trial | **F(1,35)=4.700**<br>**p=0.037**<br>**$\eta^2_p=0.118$** | F(1,35)=0.006<br>p=0.941<br>$\eta^2_p=1.59e-04$ | F(1,35)=0.567<br>p=0.456<br>$\eta^2_p=0.016$ | **F(1,35)=9.925**<br>**p=0.003**<br>**$\eta^2_p=0.221$** |
| Physical State<br>* Trial * Group | F(1,35)=0.166<br>p=0.686<br>$\eta^2_p=0.005$ | F(1,35)=0.004<br>p=0.95<br>$\eta^2_p=1.15e-04$ | F(1,35)=0.214<br>p=0.647<br>$\eta^2_p=0.006$ | F(1,35)=.868<br>p=0.358<br>$\eta^2_p=0.024$ |

### Cognitive Task Performance

Mean response times (RTs), d’ values, and walking-*minus*-sitting differences were calculated for each participant and are plotted in **Figure 2**. Dual-task effects on each participant’s performance were categorized as improvement (higher d’ or faster response time during walking), maintenance (no change), or cost (lower d’ or slower response time during walking). The effects of age and walking on response time and accuracy (d’) were compared across trial types using 2 (trial type) x 2 (physical state) x 2 (age group) repeated measures ANOVAs (ANOVA details, F statistics, and effect sizes are presented in Table 1). The trial type factor had 2 levels as separate ANOVAs were used to interrogate mixing costs (pure vs. mixed repeat trials) and switch costs (mixed repeat vs. switch trials).

**Figure 2.**
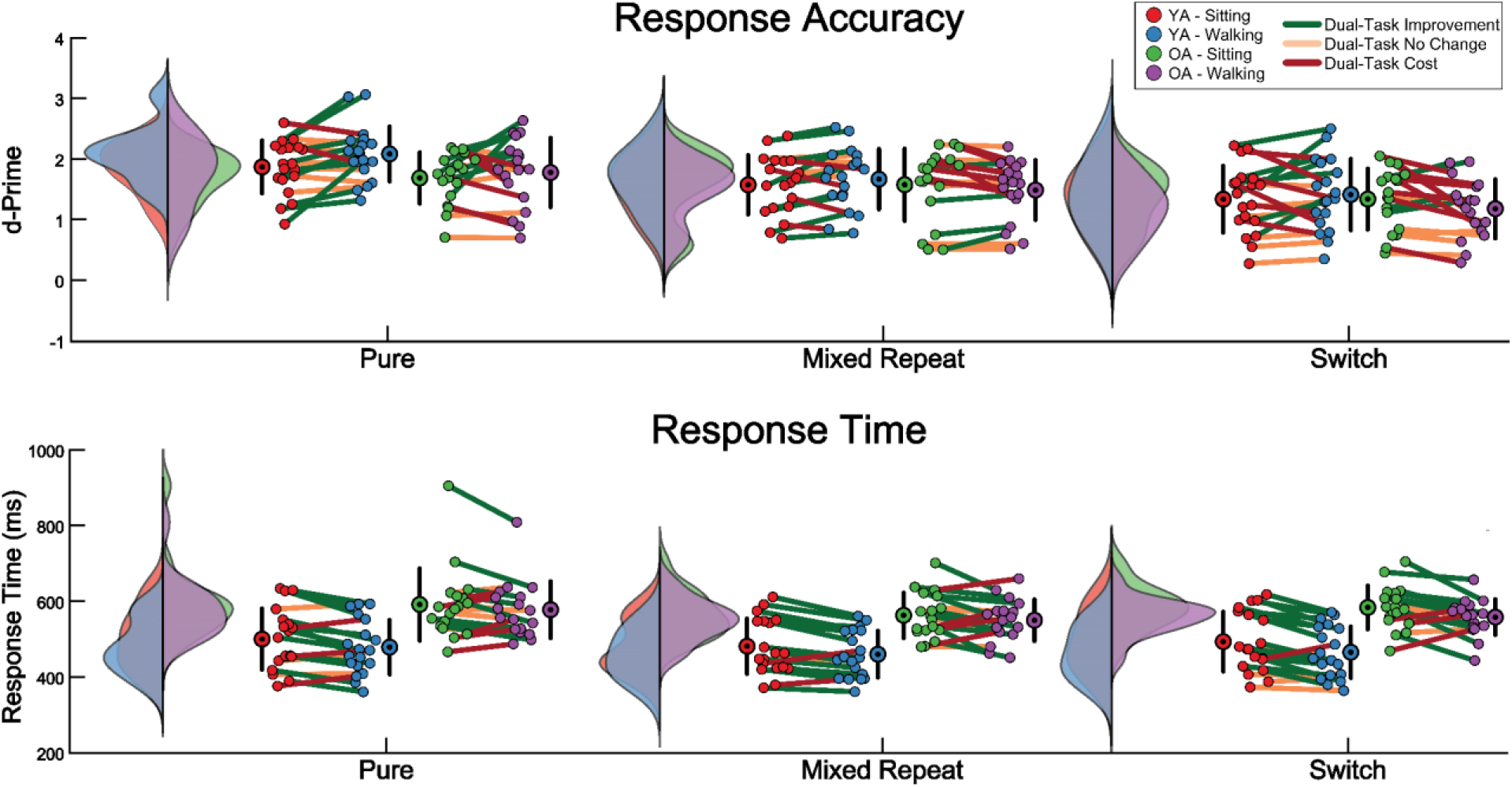
Task performance data for Younger (YA) and Older Adults (OA). Group distribution curves and individual participant values are illustrated for each condition. Dots and whiskers indicate group means +/- 1SD. Dual-Task effects (improvement, no change, or cost) are represented by the color of the line connecting each participant’s sitting and walking performance values. ***Top Panel***: d’ values. ***Bottom Panel***: response times.

### Walking Magnifies Mixing Costs

Participants must repeat the task of the preceding trial on both pure and mixed repeat trials but resolving interference between multiple tasks during mixed blocks is expected to degrade performance. Walking further increases demands. Comparing task performance on pure and mixed repeat trials revealed both response time and accuracy were affected by physical state (RT: *p=*1.655e-04; Accuracy: *p=*0.045) and trial type (RT: *p=*0.003; Accuracy: *p=*2.232e-06). A main effect of age group on RT (*p=*2.492e-04) and interaction effects on accuracy between physical state and trial type (*p=*0.037) and physical state and age group (*p=*0.047) were also observed (Table 1, columns 1 and 2).

Accuracy was higher on pure than mixed repeat trials when participants were sitting (*p=*0.0034, d=0.407), and walking (*p=*2.353e-07, d=0.708), but the mixing cost increased when participants walked. The larger cost reflected paradoxical dual-task improvement to accuracy on pure trials (*p=*0.0211, d=0.306), but no change to mixed repeat accuracy (*p=*0.9, d=0.005). This widened the gap in accuracy between trial types when participants walked. At the group level, dual-task improvement was limited to younger adults (*p=*0.0053, d=.309). Collectively, older adults’ accuracy on pure and mixed repeat trials neither improved, nor worsened when they walked at the same time (*p=*0.9, d=0.002). However, the colored lines in **Figure 2** illustrate that all three possible dual-task accuracy outcomes (improvement, no change, and cost) were represented by individuals from each age group. Younger adults overall were not more accurate than older adults, but they responded substantially faster (*p=*0.0002, d=1.244). RTs for both age groups were faster during walking (*p=*0.0002, d=0.238). Contrary to expectations, RTs were faster on mixed repeat trials than pure trials (*p=*0.0027, d=0.323).

### Switch Costs: Older Adult Performance Deteriorates As Cognitive-Motor Demands Increase

In mixed blocks, switch costs to accuracy and RTs are thought to reflect the additional processing time required for disengagement from the previous trial’s task, and activation of a different task. Walking while switching between tasks represents the greatest cognitive-motor load participants experience in the current experimental paradigm. Comparing task performance across mixed block trials revealed accuracy depended on the trial type (*p=*3.484e-07), and that there was an interaction effect between physical state and age group (*p=*0.008). There were also main effects of physical state (*p=*1.178e-04), trial type (*p=*0.003), age group (*p=*8.898e-05), and an interaction effect between physical state and trial type (*p=*0.001) on participants’ response times (Table 1, columns 3 and 4). As expected, accuracy was lower on switch trials (*p=*3.485e-07, d=0.492). Younger adults’ accuracy on mixed repeat and switch trials did not change during walking (*p=*0.1181, d=0.161), but older adults’ accuracy decreased (*p=*0.0294, d=0.234). Individual participant differences demonstrate accuracy costs during walking were more pronounced in some than others. Not every older adult suffered dual-task performance costs, and not every younger adult improved. 33.3%, 44.4%, and 50% of older adults suffered dual-task costs to accuracy on pure, mixed repeat, and switch trials respectively. In comparison 10.5%, 21.1%, and 31.6% of younger adults suffered dual task accuracy costs on pure, mixed repeat, and switch trials respectively. Response times were slower on switch trials when participants sat (*p=*5.330e-05, d=0.258), but this switch cost was not statistically detectable during walking (*p=*0.0503, d=0.106). Responses to both mixed repeat (*p=*0.0013, d=0.268) and switch trials (*p=*3.595e-05 d= 0.420) became faster during walking, but the effect was larger on switch trials (Fig. 2). As above, younger adults responded considerably faster than older adults (*p=*8.898e-05, d=1.393).

### Task Performance Summary

When the task was less challenging (pure blocks), younger adults responded faster and more accurately when walking, suggesting performance was enhanced by the motor load. As the cognitive task became more difficult, younger adults continued to respond faster when walking, while also maintaining the same levels of accuracy as when seated. Older adults also responded faster when walking, but their accuracy decayed if the trial was difficult. This may reflect more flexible resource allocation at higher cognitive-motor demands by younger adults than older adults. Faster response times may reflect a change in strategy to release cognitive resources by ending trials quickly, notwithstanding accuracy costs incurred by older adults.

Switch costs to response times disappeared when subjects walked. This could indicate participants improved the efficiency of switch-specific processes. Alternatively, it could indicate participants became indiscriminate in their handling of mixed block trials when forced to walk at the same time. It is possible the underlying explanation varied from one participant to another.

### Gait Kinematics

Dual-task kinematic measurements normalized to baseline gait are presented in **Figure 3**. The effects of age and performing the cognitive task on gait characteristics were assessed using 3 (block type) x 2 (age group) repeated measures ANOVAs (details, F statistics, and effect sizes are presented in Table 2). There was a main effect of block type on stride time (*p=*7.375e-04) and stride time variability (*p=*0.001), and an effect of age group on stride length (*p=*3.636e-06), stride length variability (*p=*1.441e-05), stride time (*p=*5.743e-06), and stride time variability (*p=*4.910e-05). There was also an interaction between block type and age group on stride time (*p=*0.005) (Table 2).

**Figure 3.**
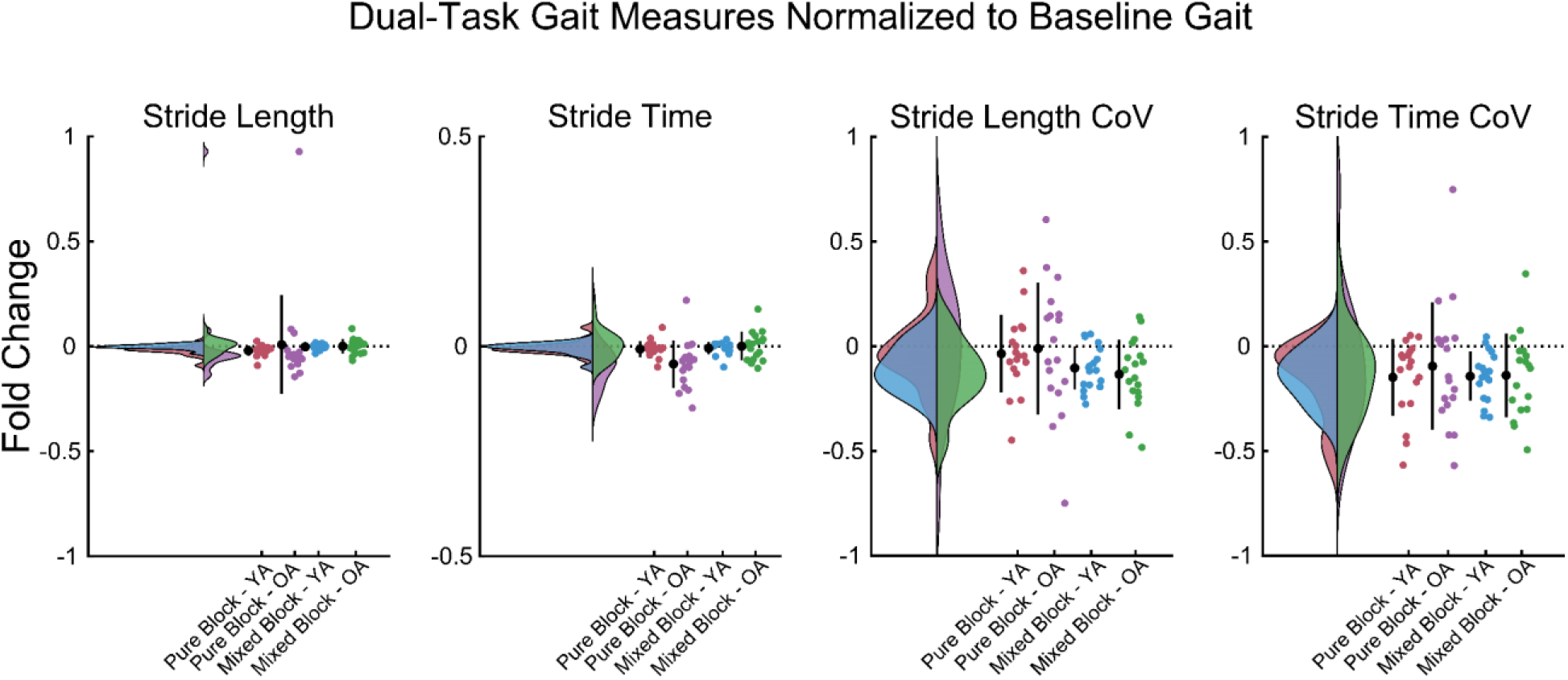
Kinematics during dual-task walking normalized to baseline gait (walking only). Colored dots show each participant’s dual-task fold change from baseline for stride length (m), stride time (s), stride length variability (coefficient of variation), and stride time variability (coefficient of variation) during pure and mixed blocks. Group distributions are shown for each condition at the left of each plot. Black dots and whiskers indicate group means +/- 1SD.

**Table 2:**
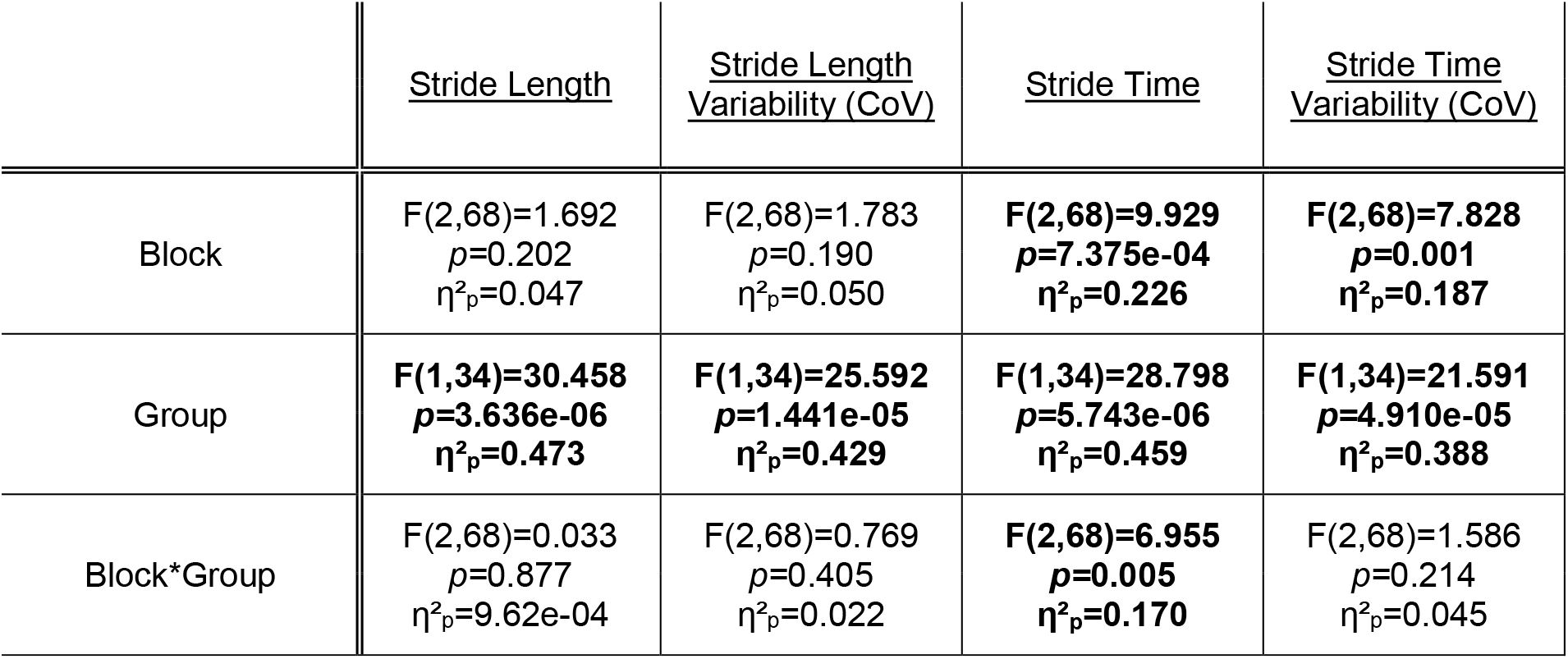
Results of repeated measures ANOVAs comparing gait measures to assess age and block (walking only, pure block, mixed block) effects. Bold text indicates *p*-value < 0.05. Effect size is reported as partial eta squared (η²_p_).

Older adults took shorter (*p=*3.636e-06, d=1.791) and slower strides (*p=*5.743e-06, d=1.748) that were less consistent in length (*p=*1.441e-05, d=1.523) and duration (*p=*4.910e-05, d=1.491) than younger adults (Fig. 3). Gait consistency of both younger and older adults improved when they performed the cognitive task simultaneously. Stride time variability for both age groups was lower in pure (*p=*0.0261, d=0.245) and mixed (*p=*7.214e-04, d=0.286) blocks than walking only. The negative normalized values and distributions for stride time variability shown in **Figure 3** illustrate how most participants’ gait became less variable as cognitive motor demands increased. Older adults took quicker strides in pure blocks than mixed (*p=*1.711e-05, d=0.437) or walking only blocks (*p=*9.928e-04, d=0.427), but there was no difference in stride duration between mixed and walking only blocks (*p=*0.988, d=0.009). This may reflect adaptation to treadmill walking, as pure blocks necessarily occur early in the experiment, or an attempt by older adults to reduce single support time (i.e. the swing phase of a step where only one foot is engaged with the ground) by taking faster steps more frequently.

### Electrophysiology Overview

Electrode channel clusters across four pre-defined intervals of proactive and reactive processes were selected to assess age-differences in neural changes occurring during walking. Clusters and intervals for amplitude measurements were defined by characterizing statistically significant walking-sitting differences in a previously published dataset of young adults performing the same task (Richardson et al., 2022).

Grand average traces and walking-minus-sitting difference topographies for late proactive processes recorded over frontal and parietal scalp are shown in **Figure 4**. Electrophysiology for reactive processes is shown in **Figures 5, 6, and 7**. Baselines were calculated over the 100 ms interval preceding cue onset (0 ms, vertical red line). Amplitude was measured relative to the baseline. It is important to note that these ERPs reflect the average neural response of anticipatory and reactive processes on trials to which a correct response was provided.

**Figure 4.**
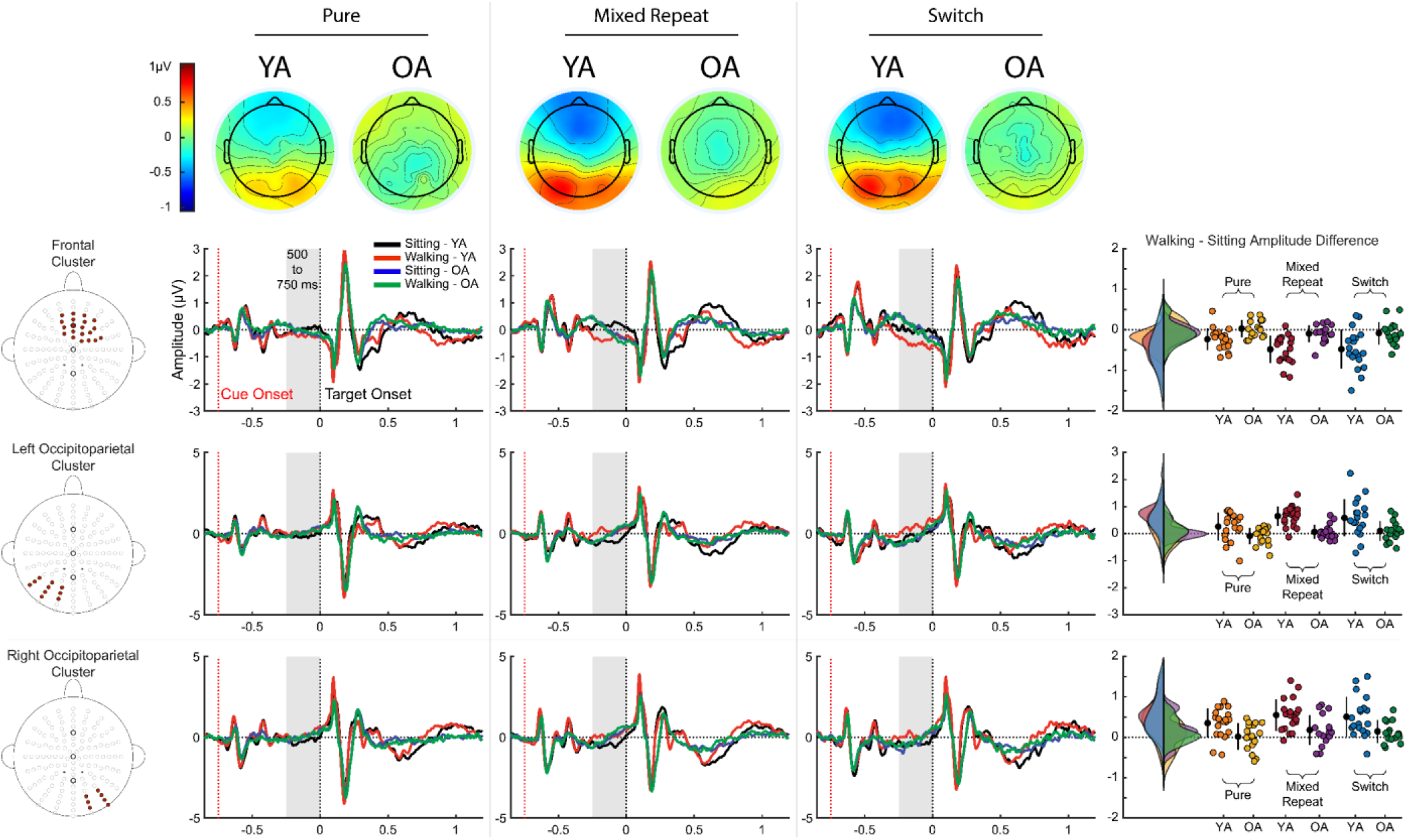
Electrophysiological data for the preplanned 500-750ms interval (late cue-target interval). Grand average ERP traces (average of channel clusters displayed to the left of each row) show younger (YA) and older adult (OA) cue and target evoked responses for the trial type listed at the top of each column (From left to right: Pure, Mixed Repeat, Switch). Raincloud and scatter plots display individual subject means (colored dots), group means +/- 1SD (black dots and whiskers), and group distribution curves for walking-*minus*-sitting amplitude differences at each channel cluster across the 500-750ms interval. Topographical maps at the top of the figure plot the walking-*minus*-sitting amplitude difference across the entire scalp.

**Figure 5.**
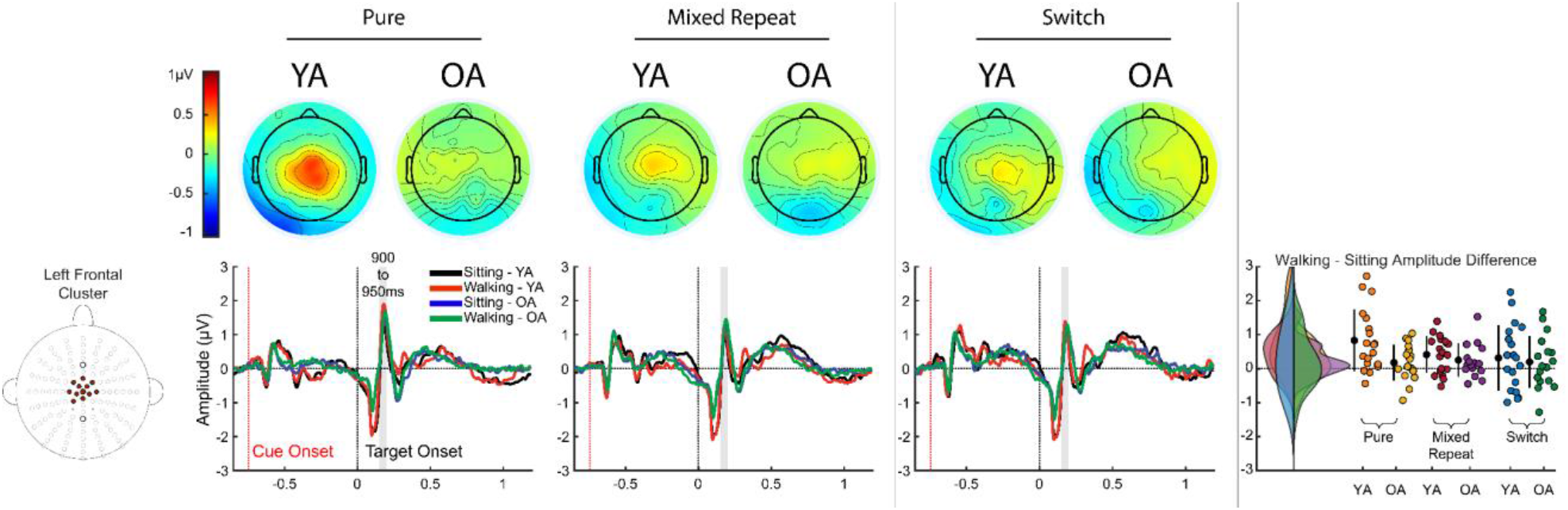
Electrophysiological data for the preplanned 900-950ms interval (150-200ms post-target). Grand average ERP traces (average of channel clusters displayed to the left of each row) show younger (YA) and older adult (OA) cue and target evoked responses for the trial type listed at the top of each column (From left to right: Pure, Mixed Repeat, Switch). Raincloud and scatter plots display individual subject means (colored dots), group means +/- 1SD (black dots and whiskers), and group distribution curves for walking-*minus*-sitting amplitude differences at each channel cluster across the 900-950ms interval. Topographical maps at the top of the figure plot the walking-*minus*-sitting amplitude difference across the entire scalp.

**Figure 6.**
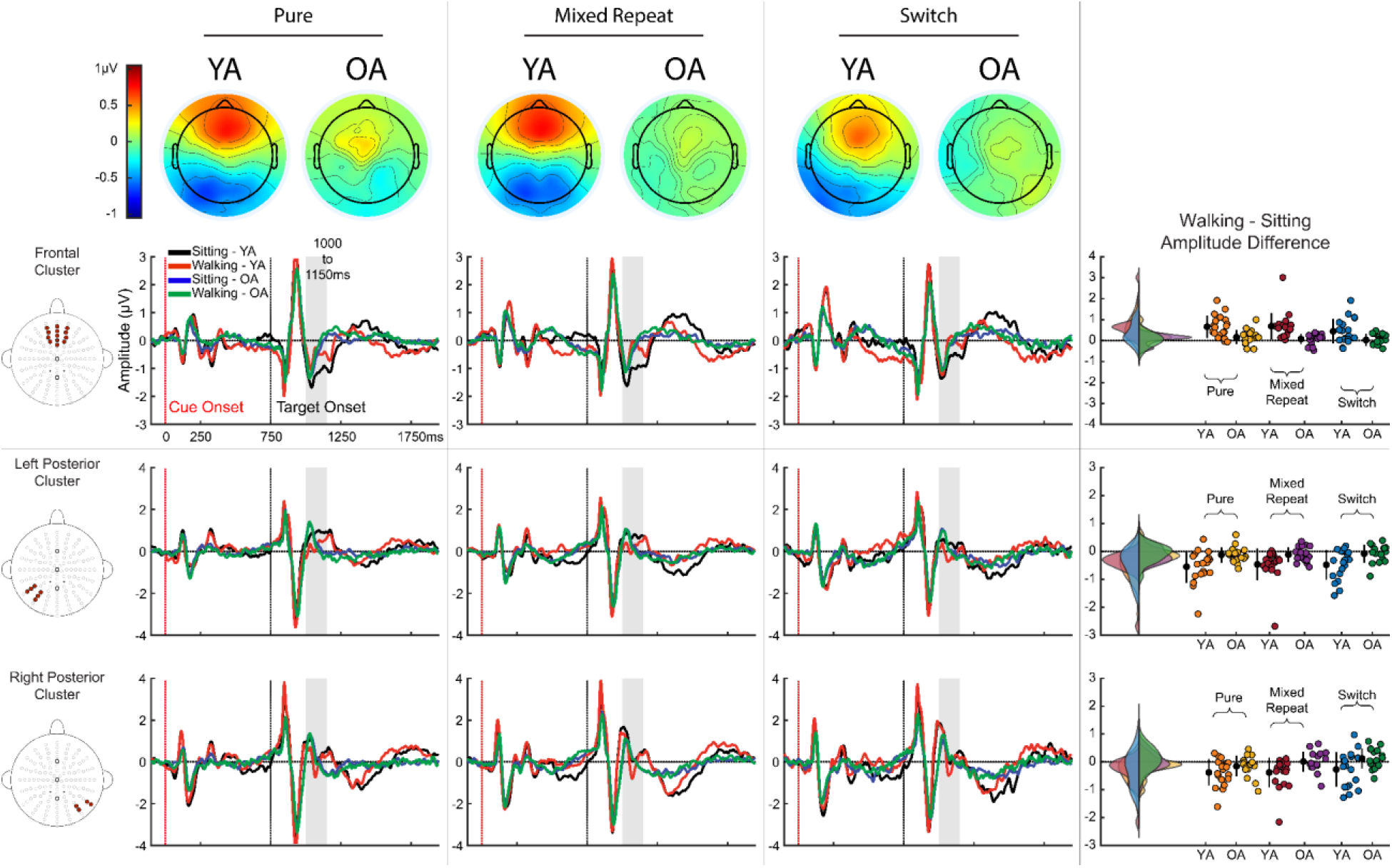
Electrophysiological data for the preplanned 1000-1150ms interval. Grand average ERP traces (average of channel clusters displayed to the left of each row) show younger (YA) and older adult (OA) cue and target evoked responses for the trial type listed at the top of each column (From left to right: Pure, Mixed Repeat, Switch). Raincloud and scatter plots display individual subject means (colored dots), group means +/- 1SD (black dots and whiskers), and group distribution curves for walking-*minus*-sitting amplitude differences at each channel cluster across the 1000-1150ms interval. Topographical maps at the top of the figure plot the walking- *minus*-sitting amplitude difference across the entire scalp.

**Figure 7.**
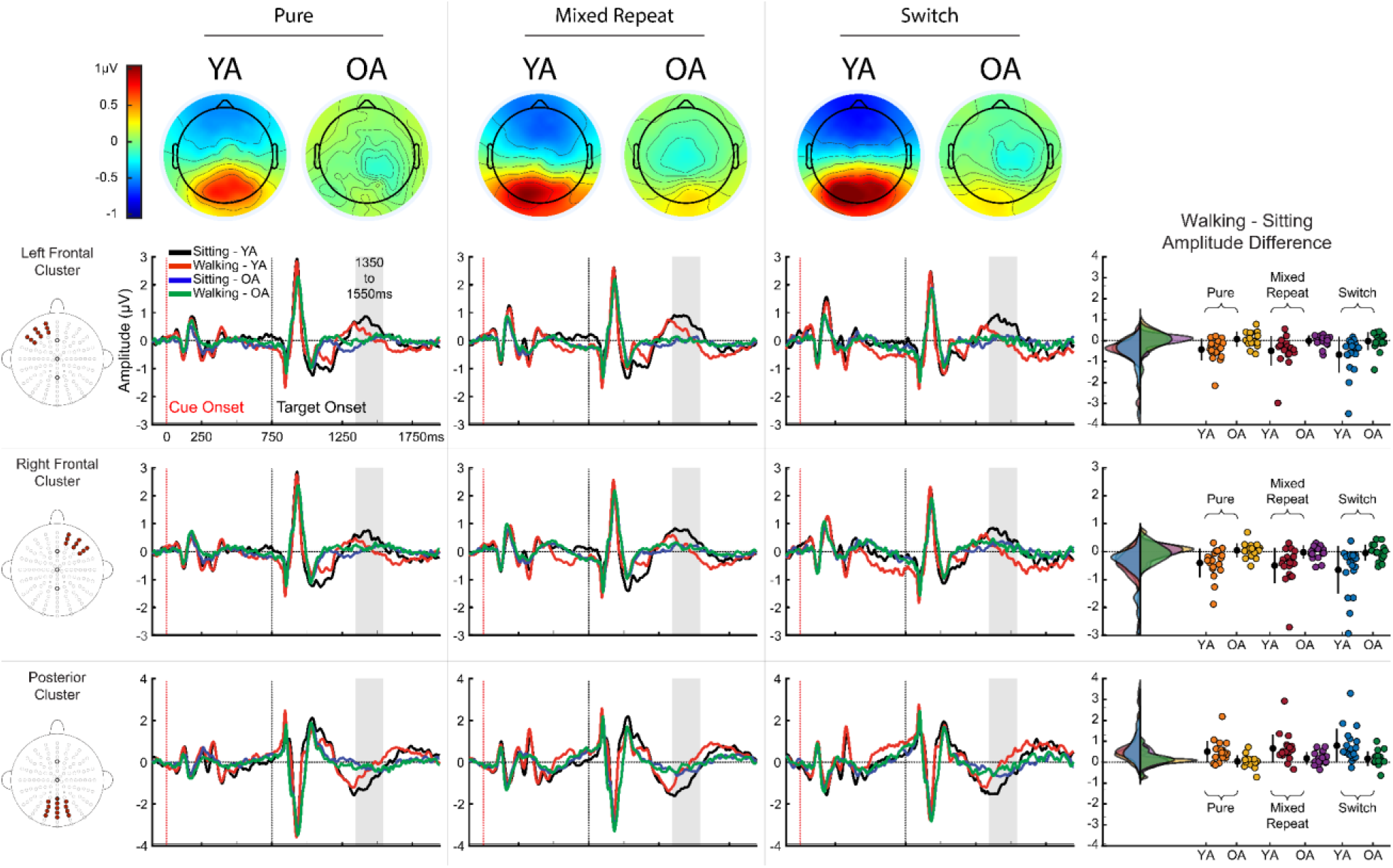
Electrophysiological data for the preplanned 1350-1550ms interval (late post- target interval). Grand average ERP traces (average of channel clusters displayed to the left of each row) show younger (YA) and older adult (OA) cue and target evoked responses for the trial type listed at the top of each column (From left to right: Pure, Mixed Repeat, Switch). Raincloud and scatter plots display individual subject means (colored dots), group means +/- 1SD (black dots and whiskers), and group distribution curves for walking-*minus*-sitting amplitude differences at each channel cluster across the 1350-1550ms interval. Topographical maps at the top of the figure plot the walking-*minus*-sitting amplitude difference across the entire scalp.

Post-hoc testing of ANOVA results focused on main effects and interactions involving physical state or age group factors to prioritize the study of age-differences in the flexibility of neural responses during walking. Other statistical effects are noted (i.e. main effects of trial type or channel cluster; interactions between trial type and channel cluster), but further interrogation is deferred for two principal reasons. Firstly, locations of the channel clusters were predefined for their suitability to capture walking-dependent recalibration of control processes, as opposed to effects dependent upon the type of trial being performed. Secondly, target-evoked responses in the present study are measured relative to a pre-cue baseline, as opposed to the common methodological practice of using a separate pre-target baseline. This avoids establishing a baseline across unresolved walking-dependent changes to late proactive processes. However, this raises the possibility for distorted interpretation of target-evoked effects dependent on trial type and/or scalp region.

### Late Proactive Control: 500-750ms Post-Cue

Figure 4 displays grand average ERPs from channel clusters over frontal and lateral parietal-occipital scalp. The raincloud plots illustrate individual and group walking-*minus*-sitting amplitude differences at each channel cluster over the final 250ms of the cue-target interval (500-750ms post-cue onset; gray band on ERP plots). Topographical maps show scalp-wide mean walking-*minus*-sitting amplitude differences at this interval. A sustained frontal negativity which is more prominent on mixed block trials can be seen ∼500ms after cue onset in the neural responses of young adults during walking, but not sitting. A negativity is also seen in older adults’ neural responses, but with a later onset (∼610-650ms) and without divergence in amplitude between sitting and walking neural activity. Over posterior scalp, a sustained negativity can be seen in young adults’ neural responses while sitting, but not walking. This posterior negativity is also present in older adults’ neural responses, but no divergence in amplitude between walking and sitting cue-evoked responses can be seen. The topographic maps illustrate the net walking-*minus*-sitting amplitude differences in young adults is a frontal negativity and posterior positivity, but little difference between the walking and sitting neurophysiologies of older adults (see topographic maps in Fig. 4). A 2 (physical state) x 3 (trial type) x 3 (channel cluster) x 2 (age group) repeated measures ANOVA (F statistics and effect sizes reported in Table 3) was performed to compare the effects of walking and trial type on late proactive processes across scalp locations and age groups. This showed that amplitude depended on three-way interactions between channel cluster, physical state, and age group (*p=*1.992e-05), cluster, trial type, and age group (*p=*0.011), and cluster, physical state, and trial type (*p=*0.009) (Table 3, column 1).

**Table 3:**
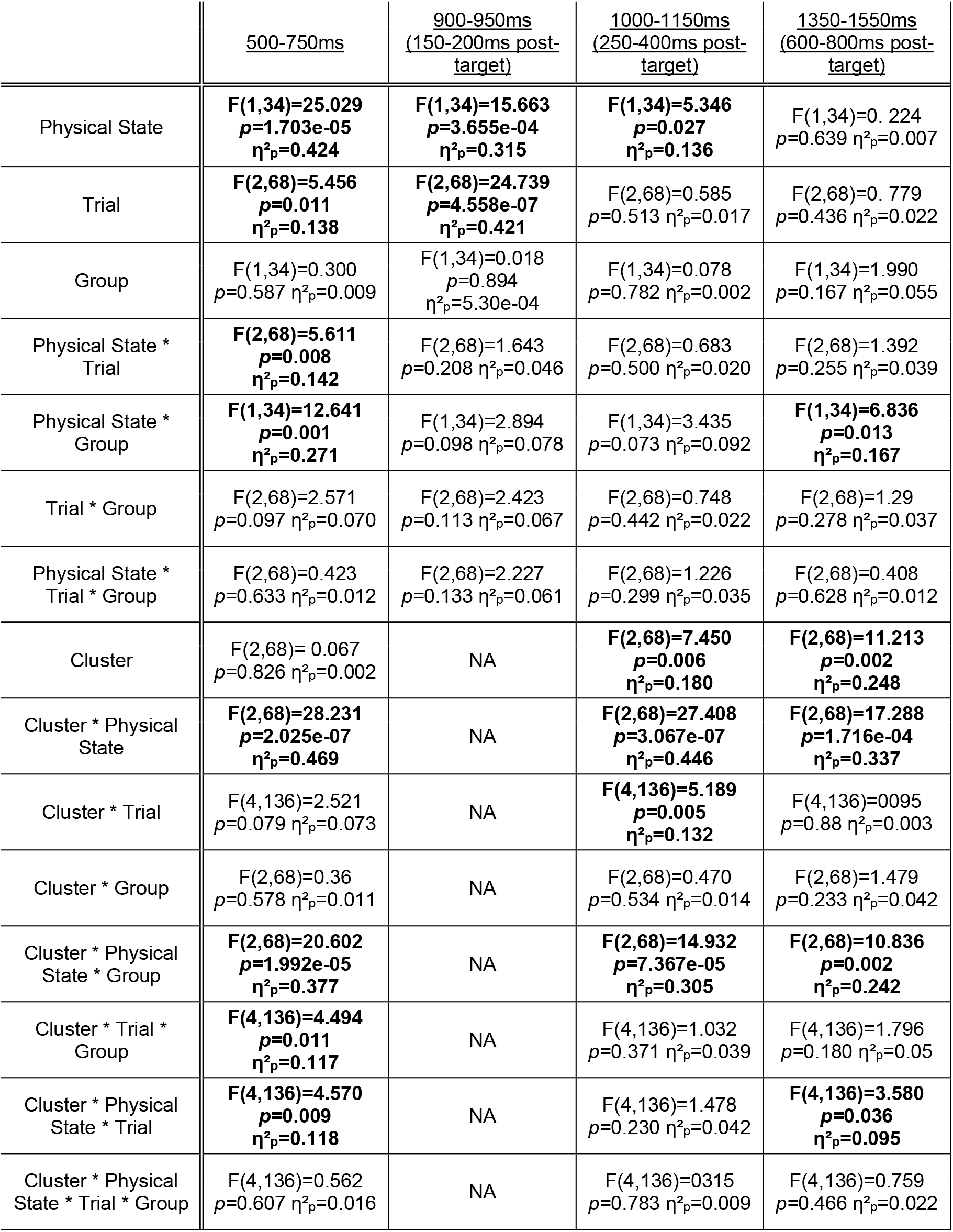
Results of repeated measures ANOVAs for ERP analysis. Bold text indicates *p*-value < 0.05. Greenhouse Geisser corrected *p*-values are shown where appropriate. Effect size is reported as partial eta squared (η²_p_).

Proactive neural activity over the frontal (*p=*8.903e-08, d=0.622), left parietal-occipital (*p=*4.620e-07, d=0.769), and right parietal-occipital clusters (*p=*5.737e-07, d=0.721) was altered when younger adults walked. Older adults’ late proactive processes did not differ during walking relative to sitting over frontal (*p=*0.961, d=0.069), left parietal-occipital (*p=*0.999, d=0.042), or right parietal-occipital scalp (*p=*0.597, d= 0.174). Absence of walking-*minus*-sitting differences may reflect older adults having failed to allocate additional resources to their preparations as flexibly as younger adults when demands increased. Late proactive processes preceding each type of trial underwent walking-dependent reconfiguration in younger adults, but the changes to the underlying neurophysiology had larger amplitudes during the more cognitively demanding mixed blocks (mixed repeat and switch trials). Walking accentuated the negativity over frontal scalp during the late preparations for pure trials (*p=*0.02, d=0.154), and this effect was even larger during mixed repeat (*p=*1.1089e-06, d=0.444), and switch trials (*p=*1.894e-04, d=0.438). Progressive walking-dependent effects on proactive processes were also measured posteriorly. During the less challenging preparations required for pure trials, walking did not change the amplitude of the underlying neural activity over left (*p=*0.217, d=0.139), but attenuated a negativity over right parietal-occipital scalp (*p=*0.003, d=0.281). When demands were higher during preparations for mixed block trials, walking-dependent changes to posterior proactive processes were more pronounced. The negativity recorded at the left and right parietal-occipital clusters was attenuated by walking during mixed repeat (left: *p=*1.104e-07, d=0.551; right: p1.969e-06, d=0.562) and switch trials (left: *p=*8.218e-04, d=0.527; right *p=*3.497e-05, d=0.501). This progression may reflect a greater requirement for walking-dependent recalibration of proactive processes during cognitive tasks which are more demanding.

### First Reactive Control Interval: 150-200ms Post-Target (900-950ms Post-Cue)

Grand average ERPs from a single centrally located channel cluster, raincloud, and topographical plots of walking-sitting amplitude differences across 900-950ms post-cue onset (150-200ms post-target) are presented in Figure 5. A divergence in amplitude between walking and sitting target-evoked responses can be seen during the fronto-centrally maximal positive peak at this interval (∼180-190ms after target onset/930-940ms after cue onset; gray band). A 2 (physical state) x 3 (trial type) x 2 (age group) repeated measures ANOVA showed that the amplitude of the positivity depended on the trial type being performed (*p=*4.558e-07) and whether the participant was sitting or walking (*p=*3.655e-04) (Table 3, column 2). Topographical plots illustrate more extreme walking-*minus*-sitting amplitude difference values at this interval when comparing young adult target evoked responses during pure trials, but there was no statistically significant interaction between physical state, trial type, and age group (*p=*0.133). The raincloud plot shows these extreme values are driven by a few individuals, which may suggest an underlying heterogeneity in the composition of the group despite similarity in age and overall neurological health.

Post-hoc analysis across both age groups showed amplification of the positive peak during trials performed while walking (*p=*3.655e-04, d=0.224). The positive peak was also larger during pure trials than either mixed repeat (*p=*1.988e-06, d=0.330), or switch trials (*p=*7.270e-05, d=0.315). There was no amplitude difference between mixed block trial types (*p=*0.9, d=0.015). Positive values in the raincloud plots illustrate the amplified positive peaks which occur during walking, but also show diversity in the responses of individuals from both age groups. Most participants demonstrate walking-related accentuation of the positive peak, but neural responses of some show the opposite: amplitude reduction during walking. These results suggest both younger and older adults recalibrate underlying neural activity of this early reactive process in response to increased cognitive-motor demands, but individual responses to walking are not uniform.

### Second Reactive Control Interval: 250-400ms Post-Target (1000-1150ms Post-Cue)

Figure 6 presents grand average ERPs from frontal and lateral parietal clusters, along with raincloud and topographical plots of walking-minus-sitting amplitude differences across 1000-1150ms post-cue (250-400ms post-target). Sustained divergences between the amplitudes of younger adult walking and sitting task-evoked activity persist across a frontal negativity, and a positivity seen at both posterolateral channel clusters (gray band). No such sustained differences can be seen when comparing the walking and sitting target-evoked responses of older adults. A 2×3×3×2 repeated measures ANOVA showed amplitude of neural activity underlying the reactive processes at this interval depended on a two-way interaction of channel cluster and trial type (*p=*0.005) and a three-way interaction between channel cluster, physical state, and age group (*p=*7.367e-05) (Table 3, column 3).

Post-hoc tests showed that in young adults walking attenuated the negativity over frontal scalp (*p=*1.515e-07, d=0.533), the positivity over left parietal scalp (*p=*8.669e-06, d=0.443), and the positivity over right parietal scalp (*p=*5.789e-04, d=0.305). The topographical maps in Figure 6 illustrate the spatial extent of the walking-related changes to frontal and posterior neural activity underlying younger adults’ reactive processes. No corresponding walking-dependent changes to older adults’ neural activity over frontal (*p=*0.897, d=0.078), left parietal (*p=*0.811, d=0.095), or right parietal scalp (*p=*0.999, d=0.007) were identified statistically. These results suggest younger adults recalibrate reactive processes governing inhibitory control and allocation of resources to the cognitive task more flexibly than older adults. However, the raincloud plots in Figure 6 again illustrate diversity in the walking-*minus*-sitting amplitude differences of individual young adults. This may indicate there is no uniform neurophysiological response to increased cognitive-motor loads, even amidst a group of healthy younger adults.

### Late Reactive Control: 600-800ms Post-Target (1350-1550ms Post-Cue)

Figure 7 presents grand average ERPs from lateral frontal and parietal channel clusters. Walking- *minus*-sitting amplitude differences in neural activity underlying late reactive processes occurring 1350- 1550ms post-cue (600-800ms post-target) are illustrated by the topographical and raincloud plots. Sustained divergences between the amplitude of younger adult walking and sitting ERP traces can be seen at this interval (gray band). This separation can be seen in the plots for each trial type, and persists across a positivity observed at both fronto-lateral clusters, and a posterior negativity. No corresponding divergences can be seen when comparing walking and sitting ERP traces recorded from older adults. A 2×3×3×2 ANOVA showed amplitude across this interval depended on two separate three-way interactions: an interaction between channel cluster, physical state, and age group (*p=*0.002) and an interaction between channel cluster, physical state, and trial type (*p=*0.036) (Table 3, column 4).

Post-hoc tests showed walking reduced the amplitudes of the positivity over left frontal (*p=*0.001, d=0.733) and right frontal scalp (*p=*3.579e-04, d=0.724), and the parietal negativity (*p=*1.153e-05, d=1.130) in younger adults. This attenuation during walking was not detected in older adults’ neural responses over left frontal (*p=*0.999, d=0.019), right frontal (*p=*0.999, d=0.013), or parietal scalp (*p=*0.876, d=0.220). Walking-dependent reductions in the amplitude of younger adult reactive processes were detected on each trial type, but progressively increased as trials became more demanding. Walking attenuated the neural activity recorded from the left frontal cluster on pure trials (*p=*0.029, d=0.243), mixed repeat (*p=*0.011, d=0.339), and switch trials (*p=*0.005, d=0.488), but the effect was larger on mixed repeat than pure trials, and larger still on switch trials. At the right frontal cluster, walking attenuated the positivity on pure (*p=*0.013, d=0.252), mixed repeat (*p=*0.003, d=0.365), and switch trials (*p=*0.003, d=0.488), with the largest effects again occurring on switch trials. A similar sequence was observed over parietal scalp. Walking reduced the parietal negativity on pure (*p=*6.257e-04, d=0.471), mixed repeat (*p=*4.789e-05, d=0.724), and switch trials (*p=*8.697e-05, d=0.829), with the largest effects occurring on switch trials. The topographical maps in Figure 7 illustrate the extent of the frontal and parietal scalp regions over which walking attenuated the amplitude of younger adults’ late reactive neural processes. Comparing the topographical maps of different trial types demonstrates the progressive attenuation as cognitive-motor demands increase. These findings may reflect younger adults recalibrating late reactive processes with greater flexibility than older adults to match an array of cognitive-motor demands.

## DISCUSSION

### Summary of Results

The cued task-switching paradigm used in this study dissociated anticipatory processes evoked by prospective task information from reactive processes linked to target selection and response generation. Comparison of dual-task effects on task-evoked neural activity revealed age-related differences in how proactive and reactive control processes responded to cognitive-motor demands. When cognitive demands were relatively low during pure blocks, walking facilitated response accuracy in younger, but not older adults. As the task became more difficult in mixed blocks, younger adults, as a group, achieved the same level of accuracy while walking as they did while sitting, but older adults’ accuracy deteriorated. Walking progressively altered younger adults’ neural control processes as cognitive-motor demands increased. The amplitude of these walking evoked neurophysiological modulations was attenuated in the neural responses of older adults’ relative to those measured in younger adults. Lower amplitude alterations to anticipatory and reactive processes, coupled with deteriorating behavioral performance as demands increased, may reflect reduced cognitive flexibility in older adults. These results replicated, and expanded upon the findings of our previous studies (Malcolm et al., 2015; Patelaki, Foxe, Mantel, et al., 2023; Patelaki et al., 2022; Patelaki, Foxe, McFerren, & Freedman, 2023; Richardson et al., 2022).

### Greater Dual-Task Costs to Older Adults’ Cognitive Performance

Responses became faster and more accurate in younger adults when they walked during pure blocks. One possible explanation for this boost in performance is an elevated state of arousal brought on by the physiological response to walking (Schaefer, Lövdén, Wieckhorst, & Lindenberger, 2010). The benefits of walking to younger adults’ accuracy were not without limits, however, and dissipated when the task became more challenging. Older adults fared worse. Their accuracy decayed when they walked during more difficult task blocks, although they continued to provide faster responses than they had while sitting. The variable impact walking had on cognitive performance suggests changes in the difficulty of the cognitive task shifts the balance between dual-task benefits and costs.

Faster response times during dual-task walking could also reflect a strategic compromise to limit overlap between cognitive and motor demands (Tomporowski & Audiffren, 2014). Providing faster responses may resolve reactive control processes sooner, relieving interference to motor processes stemming from the cognitive task. Older participants continued to provide faster responses as cognitive-motor demands increased, despite the dual-task costs to accuracy. Switch costs to response time also disappeared when participants walked while performing the task. Delayed responses during switch trials reflect additional time taken to disengage from one task, and activate another (Tomporowski & Audiffren, 2014; Whitson et al., 2014). This constellation of lower accuracy, faster response times, and erasure of response time switch costs may indicate older adults became more haphazard and accepted a speed-accuracy tradeoff to relieve cognitive-motor demands. As no specific prioritization instructions were provided to participants, this may reflect measures to safeguard gait stability under dual-task conditions.

Response times were unexpectedly faster during mixed repeat trials compared to pure trials, the opposite of what is typically predicted in task-switching experiments. Mixing costs to response time and accuracy are thought to reflect interference stemming from the task which is not currently being performed, but must be maintained in working memory (Bugg & Braver, 2016; P. S. Cooper, Garrett, et al., 2015; F. Karayanidis, Whitson, et al., 2011; Tarantino, Mazzonetto, & Vallesi, 2016; Whitson et al., 2014) and to persistence of activity in task-relevant perceptual processing regions of cortex involved in the to-be-ignored former task (Wylie et al., 2004a). These effects are often amplified by multivalent target stimuli possessing features related to each task held in working memory (Philipp, Kalinich, Koch, & Schubotz, 2008). Despite the use of a bivalent stimulus here, response times were faster on mixed repeat than pure trials, even when participants were sitting. Mixed blocks necessarily occur after completion of all pure blocks in the current experiment. This block sequencing controls onset of task mixing effects. As stated above, the heightened state of arousal in response to walking can enhance cognitive performance. The duration of these physiological stimulating effects is unclear, but likely linger following walking cessation. Persistent states of elevated arousal could benefit subsequent blocks, and contribute to the unusually fast mixed repeat trial response times seen here. Accuracy did suffer mixing costs, but nonetheless may have also benefitted from persisting effects of walking. Task strategy on mixed repeat trials may also be impacted by adaptations which participants carried forward after the earliest dual-task blocks. Completing a version of this experiment with a single physical state may clarify the local (within block) and lingering (between blocks) effects of walking.

Several older adults experienced dual-task accuracy improvement when demands were low, and not all suffered dual-task costs when demands were high. Dual-task effects were also heterogeneous in healthy younger adults, a group presumed to possess superior capacity (Patelaki et al., 2022; Richardson et al., 2022). A complex interaction between physiological, pathological, and even elective factors like task prioritization may define the threshold level of demands which separates adaptation to dual-task walking, and performance decay. The threshold may also change from one moment to the next, a dynamic operation which individuals possess limited agency over (Fraser et al., 2016; Varela-Vasquez, Minobes-Molina, & Jerez-Roig, 2020). Using this threshold to stratify groups into dual-task improvers and non-improvers effectively differentiates between neural responses which are and are not accentuated by dual-task walking (Patelaki et al., 2022). The lower ratio of non-improvers to improvers among the younger adults may partly explain the age-differences in dual-task proactive and reactive neural changes. Heterogeneity also suggests the group level analysis may mask dual-task neural changes to the control processes of some older adults. Stratification of a larger group of older adults on the basis of dual-task effects on performance may be necessary to comprehensively elucidate age-related neural differences.

### Facilitation of Gait Stability by a Simultaneous Cognitive Task

The duration of both younger and older adults’ strides became more consistent when a cognitive task was performed at the same time. Reduced variability in gait metrics may reflect an overall improvement in stability, and a commensurate reduction in fall risk (Nakamura, Meguro, & Sasaki, 1996). Facilitation of gait regularity by a simultaneous cognitive task may seem counterintuitive, but this effect has been repeatedly demonstrated when cognitive demands are modest (Decker et al., 2016; Hamacher et al., 2019; Patelaki et al., 2022; Richardson et al., 2022). Proactive and reactive control of motor processes subserving gait improves responsiveness to balance perturbations and navigation of hazards or environmental changes (Haefeli, Vögeli, Michel, & Dietz, 2011; Nordin, Hairston, & Ferris, 2019; Potocanac, Smulders, Pijnappels, Verschueren, & Duysens, 2015; Sipp, Gwin, Makeig, & Ferris, 2013; Wagner, Makeig, Gola, Neuper, & Müller-Putz, 2016; Wagner, Martínez-Cancino, & Makeig, 2019). Reducing the automaticity of motor processes enables tailoring gait to environmental demands, but counterproductively increases the susceptibility of motor processes to endogenous noise, which degrades consistency (Beilock et al., 2002; Beilock & Gray, 2012; Hamacher et al., 2019). Mild cognitive demands which redirect overt attention away from control of motor processes can reduce sensitivity to endogenous noise. Whether this effect is a benefit or a hindrance is context dependent. On an uneven walking surface with irregularly spaced obstacles, a highly automatized gait may be ill suited for safe navigation. Alternatively, on a flat surface with consistent features, an automatic gait pattern might be ideal, liberating attention for other endeavors. Obviously in the current paradigm, treadmill walking aligns best with this latter explanation.

The effect reverses as cognitive tasks become increasingly challenging, heightening cross-modal resource competition. The result is a “U” shaped plot of gait variability measures against simultaneous cognitive demands (Decker et al., 2016; Lövdén et al., 2008). The inflection point beyond which cognitive demands degrade gait consistency occurs at lower demands in older adults (Decker et al., 2016). The improved gait regularity observed in the current experiment may indicate cognitive-motor demands were not challenging enough to push either younger or older adults past this inflection point. One of the strengths of this paradigm is the ease with which cognitive-motor demands can by adjusted by changing the difficulty of the cognitive task, the motor task, or both. Fixed-speed treadmill walking at a preferred pace represents one of the least intensive forms of walking (Kao & Pierro, 2021; Tomporowski & Audiffren, 2014), yet elicited dual-task costs to the cognitive performance of most older adults and even some younger adults in the present work. Titrating dual-task costs to gait variability could be accomplished by increasing difficulty of the cognitive task, or escalating motor demands with non-preferred or variable walking paces, self-paced treadmill walking, or overground walking (Kao & Pierro, 2021; Tomporowski & Audiffren, 2014).

### Reduced Neurophysiological Markers of Flexibility in Older Adults

During dual-task walking younger adults as a group did not suffer performance costs, and modulated the anticipatory and reactive processes which underlie task preparation and response generation. The more challenging a trial was, the larger the amplitude of the dual-task changes were. In contrast, dual-task walking did not modulate the neural activity of older adults as extensively. Rather, both proactive and reactive neural activity was very similar while walking and sitting in older adults, when they performed the task correctly.

The amplitude of younger adults’ neural responses changed across the range of experimental conditions. Distinct proactive and reactive neurophysiological modulations were evident depending on whether younger adults were sitting, or walking, and whether they were performing a pure, mixed repeat, or a switch trial. Progressive accentuation of neural activity that depended on task difficulty was measured at time intervals corresponding to the cue-evoked centroparietal positivity and frontal negativity, as well as late target evoked reactive neural activity. The cue-evoked centroparietal positivity and frontal negativity have been associated with activity in the PPC and DLPFC respectively (S. Jamadar et al., 2010; Frini Karayanidis & Jamadar, 2014; F. Karayanidis et al., 2009). These late proactive processes reflect development and maintenance of plans as the onset of the trial looms. Switch trials tend to evoke larger amplitudes during these intervals, likely reflecting the greater anticipatory demands preceding a task switch (Falkenstein, Hoormann, Hohnsbein, & Kleinsorge, 2003; S. D. Jamadar et al., 2015; F. Karayanidis, Provost, Brown, Paton, & Heathcote, 2011; F. Karayanidis, Whitson, et al., 2011; Kray, Eppinger, & Mecklinger, 2005; Mansfield et al., 2012; West, 2004). The amplitude of the frontal negativity is also correlated with better task performance (Gajewski et al., 2010). Progressive amplification of these neural processes during dual-task walking may reflect additional recruitment of the DLPFC and PPC as cognitive control is marshaled to maintain cognitive performance in younger adults. The progressive effects to the late reactive processes have less clear associations, and have not been discussed as extensively in the extant task-switching literature. These dual-task neural changes possibly reflect earlier resolution of the reactive control processes, and a more rapid return to baseline at the end of a trial. This may be advantageous from the perspective of brain resource economy by limiting the duration of overlap between cognitive and motor processes. Dual-task walking also accentuated neural activity occurring at intervals corresponding to the target evoked P2 and N2/P3 complex in younger adults, but not in a progressive manner. These processes have been associated with activity in anterior cingulate cortex (ACC), supplementary motor area (SMA), and PPC (Iannaccone et al., 2015; Irlbacher et al., 2014; S. D. Jamadar et al., 2015), and likely reflect response selection, interference monitoring and suppression, and the allocation of cognitive resources during execution of requested cognitive tasks (Gajewski et al., 2018; Iannaccone et al., 2015; Irlbacher et al., 2014; Kok, 2001).

The dual-task neurophysiological effects identified in younger adults reproduce the results of our previous study using a completely new cohort, except for the timing of the progressive dual-task walking changes. In our previous study, progressive changes to neural activity recorded over frontal and parietal scalp were identified at the interval corresponding to the expected P3 component (Richardson et al., 2022), whereas here we see progressive frontal and parietal amplitude changes at the late proactive interval (500-750ms post-cue onset) corresponding to the cue-evoked centroparietal positivity and frontal negativity. Progressive changes to neural activity were also recorded at the late reactive interval (1350- 1550ms post-cue/600-800 post-target stimulus onset) over lateral frontal and midline parietal scalp. This discrepancy can be explained by the decision to collect all amplitude measurements relative to the pre-cue baseline period in the present study. This was an intentional methodological departure from the previous study to avoid establishing a baseline for measurement of target-evoked reactive processes across unresolved walking-dependent changes to late proactive processes.

Altering the difficulty of the cognitive task, or simultaneous motor demands, prompts changes in the neural responses of younger adults underlying task-relevant plan development and maintenance, response selection, interference monitoring, and resource allocation. These neural changes in younger adults reflect flexible cognition in response to shifts in demands. In contrast, the neural processes of older adults demonstrate lower amplitude changes in response to shifts in either cognitive or motor demands. This comparative uniformity of neural responses across diverse cognitive-motor conditions may reflect reduced cognitive flexibility in this age group.

### Study Limitations

One potential limitation of this study is that younger and older adults had different preferred walking speeds. Despite the older adults walking at slower speeds than younger adults, these paces sufficiently interfered with older adults’ performance on the cognitive task. One of the more challenging aspects of comparing cognitive-motor interactions across individuals and age groups is matching the effort a motor task requires to ensure a similar cognitive-motor load is experienced by each participant. Cognitively and physically healthy participants may still differ in many dimensions, including size, mass, adiposity, cardiovascular health, muscle mass and fiber composition, and pulmonary health. These physiological differences may contribute significantly to differences in how physically robust any two individuals who each qualify as “healthy” may be. Ensuring a consistent subjective or perceived motor load across participants will be critical to characterizing the origins of inter-individual heterogeneity in neurophysiological, kinematic, or behavioral responses to dual-task walking. Future studies may benefit by generating individualized motor loads for each participant.

Fixed-speed treadmill walking delivers a consistent motor load on a highly predictable walking surface which reduces the need to monitor for obstacles or hazards. This predictability can be an asset to experimental design and analysis, but fixed-speed walking is less demanding than self-paced treadmill walking or the overground walking used in everyday life (Kao & Pierro, 2021). Characterizing how beneficial and costly dual-task effects manifest in real-world situations will likely require repeated experimentation across different walking and environmental conditions.

### Conclusions and Future Directions

Younger adults expressed larger amplitude alterations to their neural activity than older adults in response to progressive cognitive-motor demands. The cognitive performance and gait stability of both younger and older adults may be enhanced by dual-task walking when cognitive demands are low. As demands increase, older adults increasingly show signs of dual-task costs, whereas younger adults are better able to adapt. The lower amplitude alterations to older adults’ neural responses, in combination with larger dual-task performance costs which declare sooner as demands increase, may reflect age-related reductions in cognitive flexibility. However, age-related reductions in cognitive flexibility do not necessarily represent reduced capacity. Differences in task strategy, subjective perception of fall risk or postural instability, priority differences, and non-neural physiological factors could each contribute to the observed differences in how flexibly younger and older adults respond to escalating cognitive-motor demands.

One of the barriers to comprehensively characterizing dual-task walking-related neural changes may reside in the intrinsic heterogeneity of both younger and older adults. This heterogeneity may be the outcome of a diverse set of contributing factors, including physical robustness, cardiovascular responsiveness to exercise, and voluntary factors like task prioritization. Effectively stratifying healthy age groups may be critical to the ongoing effort to relate dual-task effects across the neurophysiological, kinematic, and behavioral domains.

Providing incentive structures or cognitive, motor, or dual-task training can improve cognitive flexibility and mitigate dual-task costs (Marusic, Verghese, & Mahoney, 2018, 2022; Tait, Duckham, Milte, Main, & Daly, 2017; Wollesen, Schulz, Seydell, & Delbaere, 2017). Targeted and personalized intervention may diminish or even close age-differences in cognitive flexibility. Dual-task walking paradigms have recently gained considerable attention for their clinical utility in characterizing risk for falling and cognitive decline (M. Montero-Odasso, Muir, et al., 2012; M. Montero-Odasso, Verghese, et al., 2012; M. M. Montero-Odasso et al., 2017; Muir et al., 2012). Extending the findings of this study to understand how training, explicit priority instructions, or incentive structures alter dual-task effects to gait, cognitive performance, and the underlying neural processes, could improve the resolution of these nascent clinical assessments. Whether or not an individual benefits from interventions could be a useful tool for discriminating between people at risk for adverse health outcomes. Further characterizing the neural markers of dual-task walking, along with corresponding gait and cognitive performance costs may aid in discriminating between candidates likely to benefit from combined cognitive-physical conditioning interventions, and individuals whose dual-task costs are derived from true capacity limitations.

## Declarations

### Author Contributions

DPR, EGF, and JJF designed the study. EGF and JJF oversaw all aspects of data collection, analysis and interpretation of results. DPR collected the data. DPR analyzed the data. DPR, EGF and JJF contributed to interpreting the results and writing the manuscript.

### Consent for publication Not applicable

## Acknowledgments

We would like to thank each of the participants that enrolled in the study. Special thanks to, Nicholas Abraham, Eric Nicholas, Suzan Hoffman and Soma Mizobuchi for their help with this study.

## Ethics Statement

All procedures were approved by the University of Rochester Institutional Review Board (STUDY00001952), and complied with the ethical principles of the Declaration of Helsinki for the responsible conduct of research.

## Availability of Data and Materials

Data and custom code from this study will be made available through a public repository (e.g. Figshare) upon publication of this paper, and the authors will work with the editorial office during production to incorporate appropriate links.

## Abbreviations List

CMI: Cognitive Motor Interference
EEG: Electroencephalography
ERP: Event-related potential
MoBI: Mobile Brain-Body Imaging
CTI: Cue-Target Interval
PTI: Post-Target Interval
SMA: Supplementary Motor Area
ACC: Anterior Cingulate Cortex
PPC: Posterior Parietal Cortex

